# Climate change induced selection on and evolution of insect thermal sensitivity

**DOI:** 10.1101/2025.10.30.683704

**Authors:** Taylor M. Hatcher, J. Gwen Shlichta, Joel G. Kingsolver, Lauren B. Buckley

**Affiliations:** Department of Biology, University of Washington, Seattle, WA, USA; Department of Biology, Edmonds College, Lynnwood, WA, USA; Department of Biology, University of North Carolina, Chapel Hill, NC, USA

## Abstract

How have recent changes in temperature means and variability altered selection on and evolution of the thermal sensitivity of insect growth? We address this question for cabbage white butterflies, *Pieris rapae*, by repeating a 1999 study that linked lab measurements of the temperature sensitivity of short-term growth rates to fitness components (survival, development time, final body size, and fecundity) in the field. In 1999, selection favored increased growth at low temperatures at the expense of decreased growth at high temperatures. We document evolution consistent with this past selection: caterpillars now grow faster at low temperatures but slower at warm temperatures in 2024 compared to 1999. However, temperatures have changed rapidly, with increasing incidence of high temperatures such that selection in 2024 has reversed to favor increased growth at the highest temperature assayed. Over the 25 years, phenotypic variation has increased at warm temperatures and a tradeoff between growth at intermediate and warm temperatures has strengthened, consistent with evolutionary constraints that may restrict evolution to grow faster at rare, high temperatures. Our study demonstrates the potential for rapid evolution of thermal sensitivity in response to climate change in an agricultural pest. However, selection to initially capitalize on warming temperatures can ultimately decrease fitness as warming temperatures move into a stressful range, driving reversals in the direction of selection.

## Introduction

A pressing question posed by ongoing and accelerating climate warming is whether species can evolve their thermal sensitivity to track warming (1–5). Repeating historical laboratory and field experiments that characterize phenotypes and quantify selection on phenotypes provides an opportunity to observe the mechanisms through which organisms respond to climate change (6). Such studies will improve our limited capacity to predict varied responses to climate change (7).

Insects are important ecosystem members that are highly sensitive to environmental conditions and well documented historically (8). Dramatic declines in insect populations (9), including of butterflies in the Western US (10), reinforce the need to leverage organismal biology in predictive frameworks for ecological and evolutionary responses to climate change (11). The temperature dependence of caterpillar growth and feeding are highly relevant to fitness in the field as they determine survival, adult size and reproductive output. The temperature dependence can be quantified as thermal performance curves (TPCs) (12–14). The characteristic skewed shape of TPCs indicates evolutionary constraints that can result in tradeoffs between performances at different temperatures (e.g., specialist-generalist tradeoffs) (13, 15).

We probe the evolutionary potential of a major agricultural pest that has rapidly spread across the globe(16) by repeating an experiment on cabbage white butterflies (*Pieris rapae*) in Seattle, WA.

The experiment measured the temperature dependence of short-term growth rates in the lab before transferring caterpillars to an experimental garden to assess time and mass at pupation, survival, and egg production (17, 18). We investigate whether increases in early season temperatures and summer thermal extremes (Fig 1A) have driven selection on and evolution of thermal sensitivity. In 1999, short-term caterpillar growth rates increased with temperature up to the highest temperature assayed (35°C), which exceeded the body temperatures commonly encountered by caterpillars in summer in the field site (Fig 1B distributions). Consequently, the field study documented significant selection on growth via pupal mass to increase feeding rate at temperatures up to 23°C and to decrease feeding rate at higher temperatures (Fig 1B arrows).

**Figure 1.**
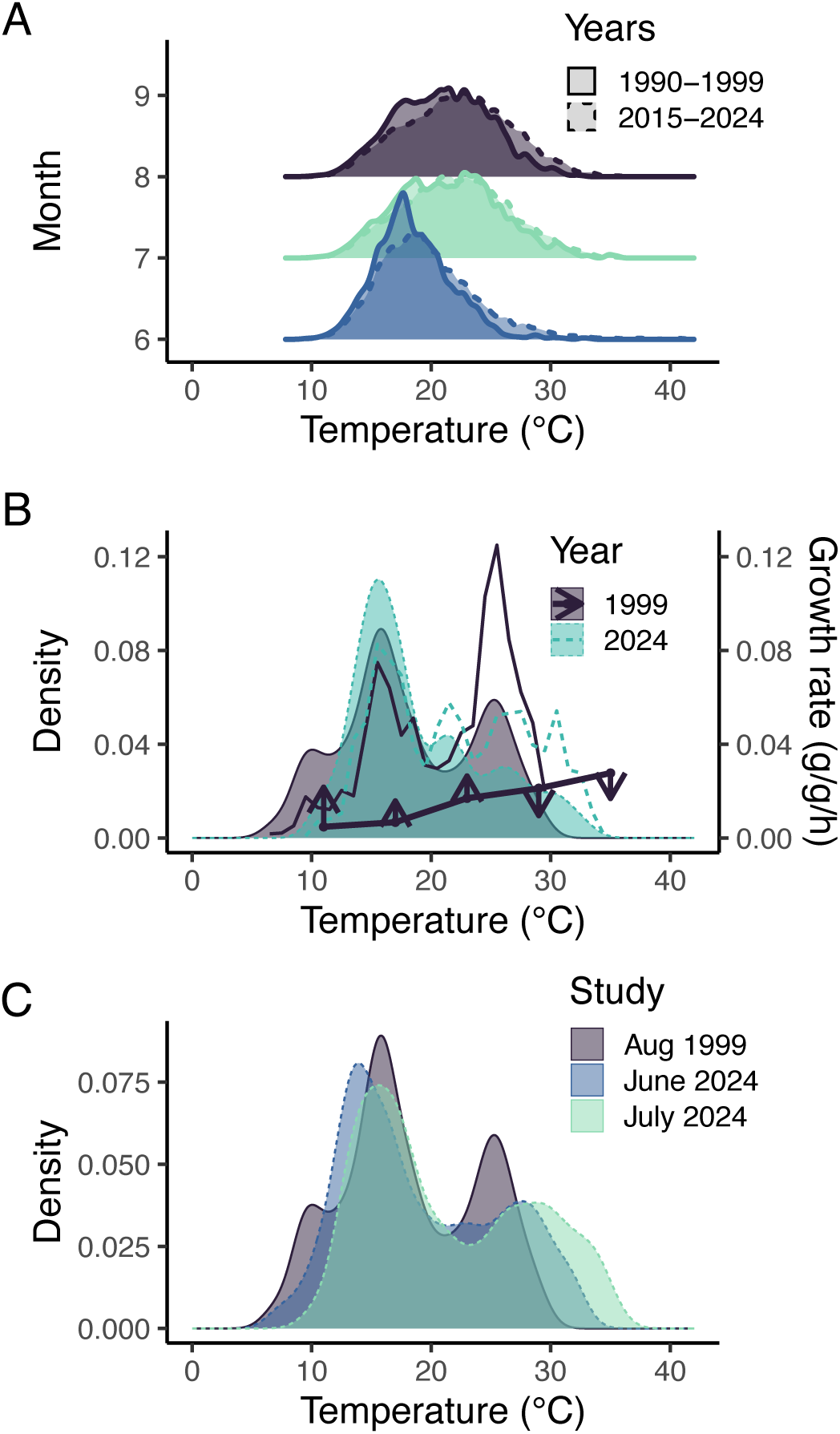
A) The distributions of monthly environmental temperatures have shifted from initial (1990-1999) to recent (2015-2024) years. B) Operative caterpillar temperatures during the dates of initial experiment (August 15-25) have increased between 1999 and 2024 (filled distributions). Growth at hot temperatures contributes disproportionately to overall growth due to higher growth rates (hollow distributions: growth rate weighted by incidence). The 1999 temperatures were cool relative the thermal optima of the experimental caterpillars’ growth, resulting in selection for improved growth at low temperatures and decreased growth at warm temperatures (line with arrows indicating the direction and strength of selection on pupal mass in the 1999 field study). The reversal in the direction of selection results from evolutionary constraints on thermal sensitivity. C) The incidence of hot operative temperatures in during the field selection studies increases from August 1999 to June 2024 to July 2024. We expect shifts in selection associated with the increased incidence of thermal extremes in 2024.

Selection via survival favored caterpillars that grew faster at 29°C. Significant selection was not observed via development time or fecundity. Selection via pupal mass across temperatures was consistent with evolutionary constraints on TPC shape: variance in growth rates was greater at higher temperatures, and a tradeoff was observed between growth rates at the two highest temperatures (29 and 35 °C). 16-43% of total phenotypic variation in growth rates was attributed to genetic variation with strong genetic correlations between high temperatures, indicating evolutionary constraints that could limit evolutionary responses (19).

We repeated the laboratory and field study 25 years later to ask whether substantial warming (Fig 1) has resulted in changes over time: Have caterpillar growth rate TPCs evolved? Have variances in and correlations among growth rates at different temperatures, which can constrain evolutionary responses, changed? Has selection on TPCs shifted? We hypothesize that temperature increases have shifted selection gradients via pupal mass and that we will find significant positive selection via survival. We anticipate TPC evolution to enable greater consumption and growth at higher temperatures, but we expect evolutionary constraints on TPC shape.

## Results

Weather station data indicates that the incidence of warm environmental temperatures over summer months have increased from the initial (1990–1999) to recent years (2015-2024, Fig 1A, ANOVA all temperatures: month F=97427.2, p<10^-15^, period F= 27984.5, p<10^-15^, month:period F= 5.0, p<0.05). Operative caterpillar temperatures during the 1999 study were cool relative to the thermal optimal for maximal growth rate (Fig 1B). Consequently, selection on pupal mass acted to increase growth rates at low temperatures at the expense of slower growth at high temperatures (Fig 1B arrows: see also Fig. 4). The same time periods during 2024 included hotter temperatures than those in 1999 (Fig 1B distributions, Welch’s t-test -5.93, p<0.001), suggesting the potential for shifts in directional selection on growth rate. Warm temperatures contribute disproportionately to growth due to faster growth rates (Fig 1B). Accordingly, the relative contributions of warm temperatures have increased from 1999 to 2024 (ANOVA temperature: F=6.2, p<0.05, period: F=1.34, p=0.2, temperature: period F=9.3, p<0.01), further suggesting the potential for shifts in selection. The incidence of warm operative temperatures in the field has increased from the August 1999 to the June 2024 to the July 2024 studies (ANOVA all temperatures: F=218.7, p<10^-15^, incidence of temperatures over 30°C: p<10^-15^, Fig 1C).

In lab assays, the temperature sensitivity of caterpillar growth rate has shifted between 1999 and 2024 toward faster growth at warm temperatures but slower growth at the hottest temperatures (Fig 2). The polynomial temperature dependence shifts significantly across time (Table 1). Mean growth rates differ significantly between 1999 and 2024 at 23°C (F[1,579]=54.6, p<0.0001) and 35°C (F[1,579]=46.4, p<0.0001; ANOVAs accounting for mother as a random effect and initial mass). The shift is consistent with evolution in response to the initially documented selection (Fig 1B).

**Figure 2.**
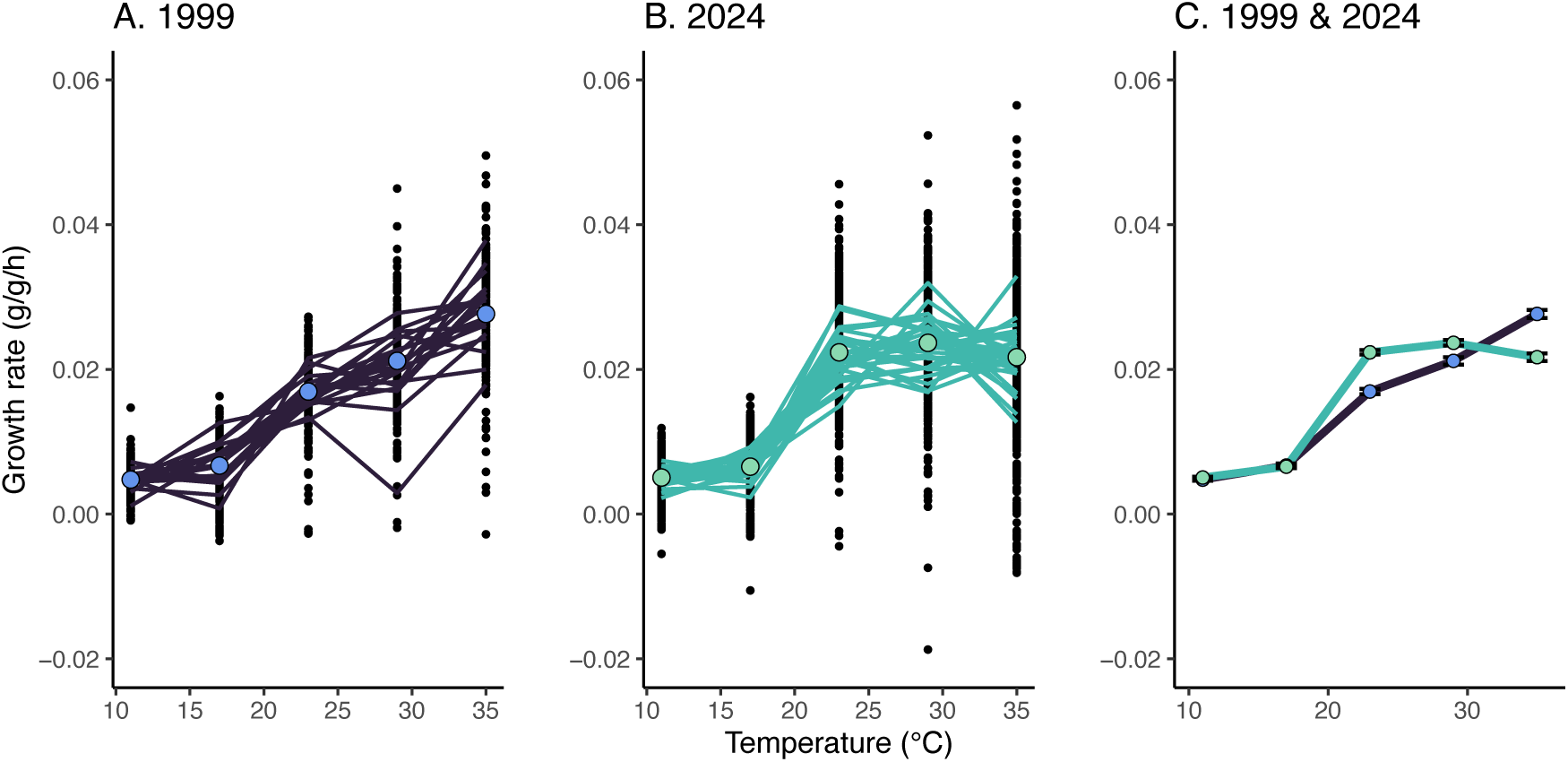
The thermal sensitivity of caterpillar relative growth rates (RGR, g/g/h) has shifted between 1999 and 2024. For A) 1999 and B) 2024, we depict individual values and means ± SE at each measurement temperature along with means by family (lines). C) Overlaying the overall means ±SE reveals growth rate increases at intermediate temperatures and declines at hot temperatures.

**Table 1.** ANOVA results from a linear mixed effects model (accounting for mother as a random effect) reveal that the temperature dependence of growth rate has changed significantly between 1999 and 2024.

|  | numDF | denDF | F | p | sig |
| --- | --- | --- | --- | --- | --- |
| **poly(temperature)** | 3 | 3187 | 1401.9 | 0 | *** |
| **year** | 1 | 3187 | 4.4 | 0.037 | * |
| **initial mass** | 1 | 3187 | 59.9 | 0 | *** |
| **poly(temp):year** | 3 | 3187 | 63.5 | 0 | *** |
| **poly(temp):initial mass** | 3 | 3187 | 6.2 | 0.0004 | *** |
| **year:initial mass** | 1 | 3187 | 3.2 | 0.0753 |  |
| **poly(temp):year:initial mass** | 3 | 3187 | 15.2 | 0 | *** |

We compared 4 recent experiments to assess whether parental or other experimental effects could account for the differences in temperature sensitivities between 1999 and 2024. Growth rates consistently decline at the hottest temperatures for each of the recent experiments (Fig S1). When including mother nested within experiment as random effects in an ANOVA (Table S1), the standard deviations of the random effects (sigma: overall=0.0085, experiment=0.0026, mother=0.0006) indicate that differences among recent experiments are substantially smaller than those between the 1999 and 2024 experiments. This points to evolution rather than parental or experimental effects as the mechanism underlying the shifts over recent decades.

Temporal shifts in both phenotypic variance covariance (random skewer correlation=0.95, p<0.0017, Fig 3A) and correlation matrices (Mantel test correlation=0.56, p<0.05, Fig 3B, Table S2) are consistent with the emergence of a tradeoff between performance at warm and hot temperatures. Variance in growth at higher temperatures has increased over time (Fig 3A). A significant negative covariation between growth rates at 29 and 35°C has emerged (interaction with year: F[1,650]=13.9, p<0.001). Eigenvectors indicating the temperatures contributing most to phenotypic variation have changed little over time: the first and second eigenvectors are dominated by variation in growth rate at 35°C and 29°C, respectively (Fig S2). The tradeoff between growth rates at 29 and 35°C also strengthens over time in the correlation matrix, which controls for the differences in variance across temperatures. In contrast, a negative correlation between growth at 11C° and intermediate temperatures has declined overtime (23°C effect of year: F[1,653]=6.2, p<0.05; 29°C effect of year: F[1,652]=5.8, p<0.05, Fig 3B). Eigenvectors indicate changing contributions to phenotypic variation at 29°C and 35°C over time (Fig S2).

**Figure 3.**
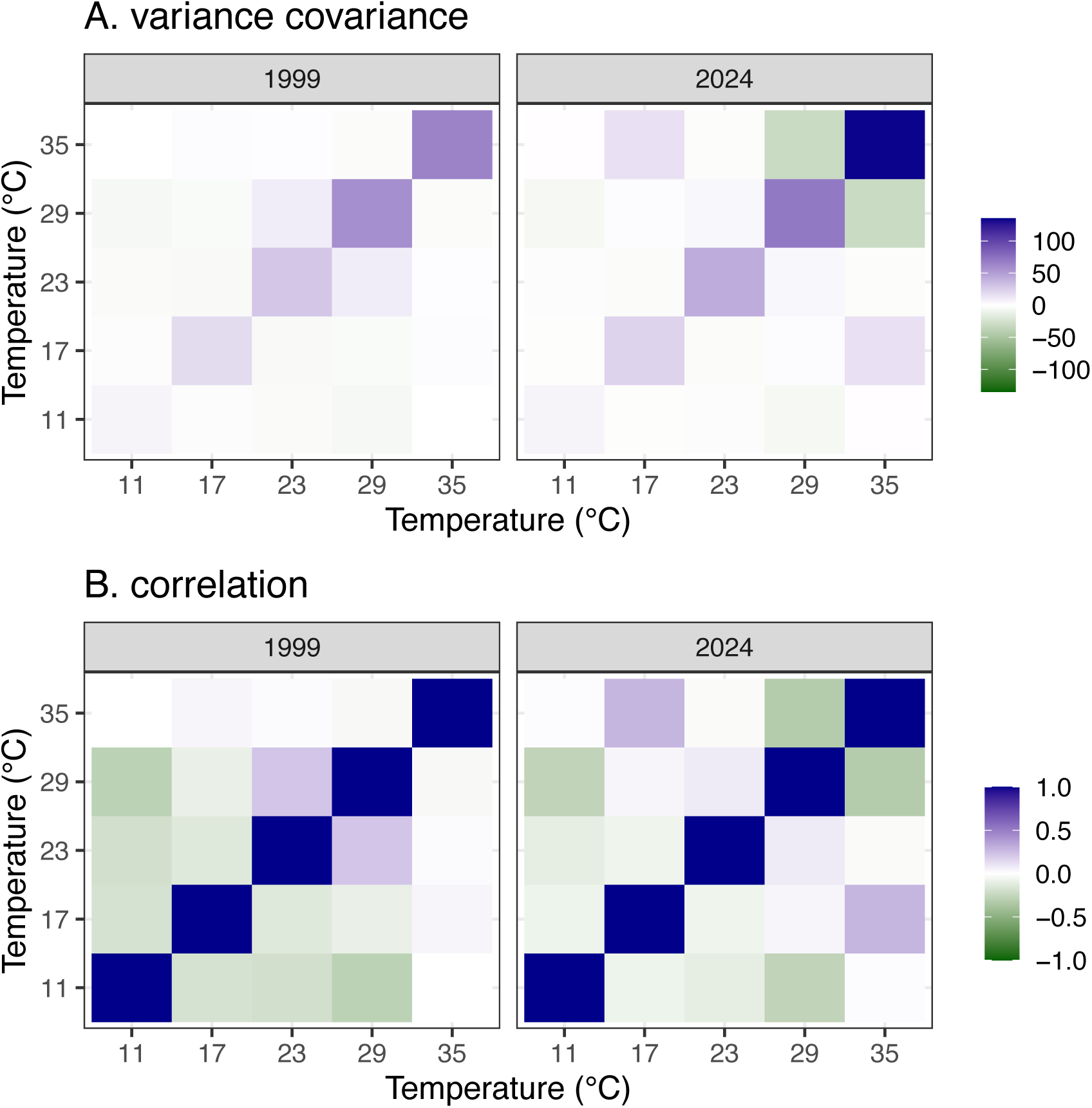
The (A) phenotypic variance-covariance and (B) correlation matrices have shifted over 25 years consistent with increasing variance in growth at high temperatures and a strengthening tradeoff between growth at 29 and 35°C.

In the field, selection on growth rate via pupal mass has reversed to favor faster growth rates at high temperatures (35°C June 2024: t[1,125]=4.5, p< 0.0001; July 2024: t[1,166]=2.5, p< 0.05, Fig 4, Fig S3, Table S3, fitness outcomes in Table S4). Positive selection via pupal mass was also observed at 23°C in June 2024 (t[1,125]=2.5, p< 0.05) and at 11°C in July 2024 (t[1,166]=2.3, p<0.05). As with the 1999 study, we found that faster growth rates at 29°C corresponded to higher survival in July 2024 (t[1,289]=2.3, p<0.05, Fig 4, Fig S4). Selection for slower development at 17°C has become significant over time (July 2024: t[1,166]=6.5, p<0.0001, Fig 4, Fig S5). We did not detect significant selection via fecundity (Figs. S6 and S7). In analyses across temperatures accounting for initial mass, the influence of growth rates at 17 and 35 °C on pupal mass differs significantly between 1999 and 2024 (Table 2). Faster growth rates at 17, 23, and 35 °C increase development rate and faster growth rates at 23 and 29°C and slower growth rates at 35 °C increase survival, but these effects do not significantly differ between 1999 and 2024 (Table 2).

**Figure 4.**
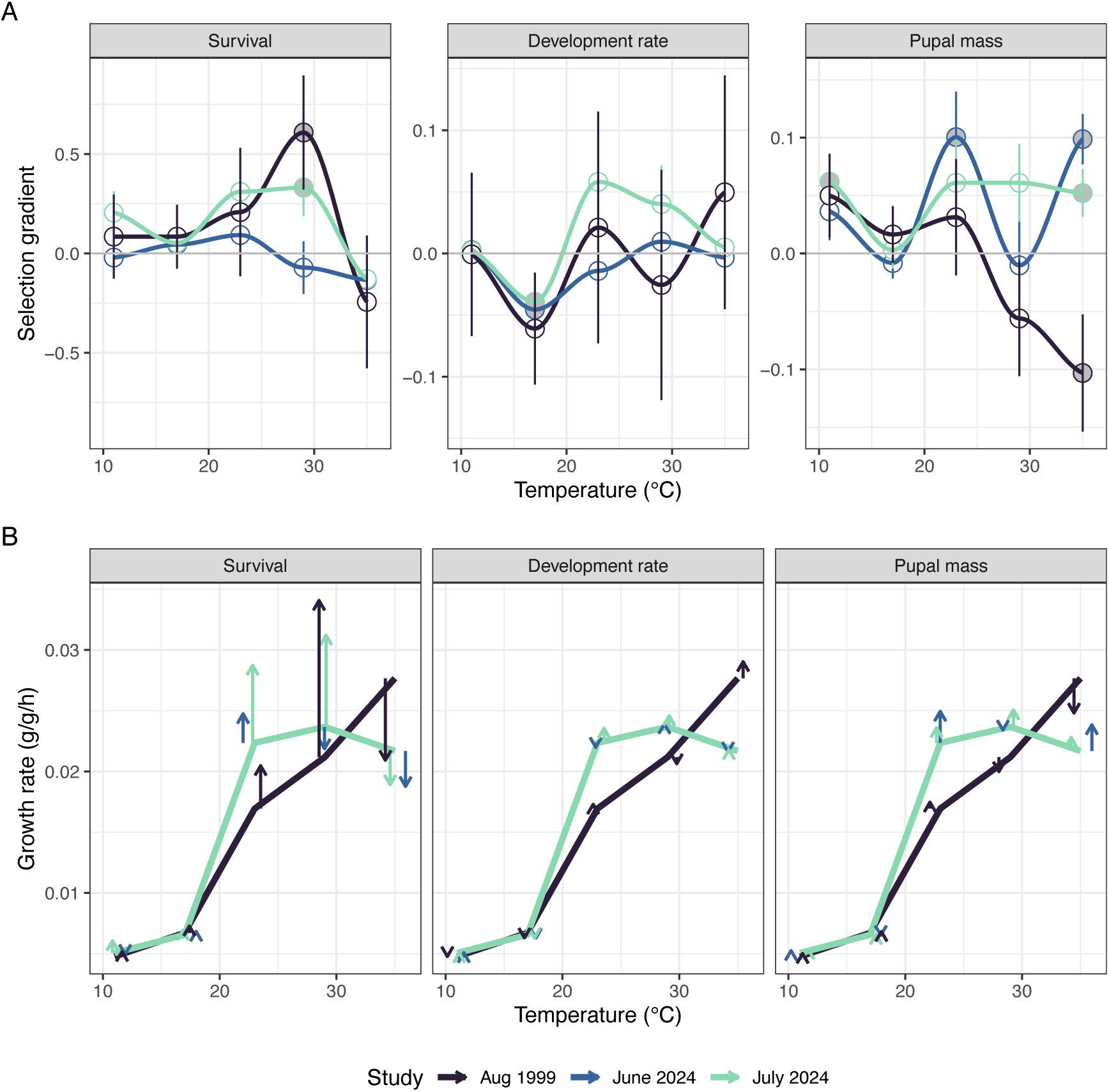
The thermal sensitivity of caterpillar relative growth rates (RGR, g/g/h) has shifted between 1999 and 2024. For A) 1999 and B) 2024, we depict individual values and means ± SE at each measurement temperature along with means by family (lines). C) Overlaying the overall means ±SE reveals growth rate increases at intermediate temperatures and declines at hot temperatures.

**Table 2.**
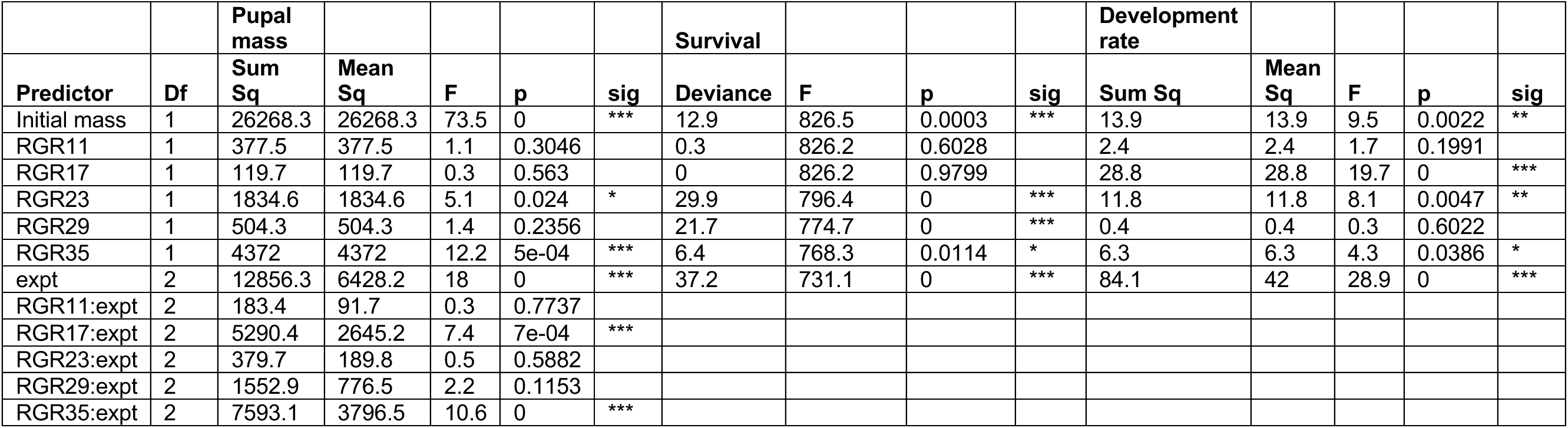
ANOVA results from (generalized) linear models examining fitness responses to initial mass, relative growth rates (RGR) across temperatures, and the three experiments. We report interactions between growth rates and experiments when the interactions received support for inclusion by Chi test comparisons.

Variation in fitness components and other traits have also shifted between the 1999 and 2024 studies. Variance in time to pupation (F[2,381]=74.7, p<10^-27^) and time to eclosion (F[2,345]=84.4, p<10^-29^) has decreased (Levene test, Fig S8). Initial mass (F[2,654]=88.7, p<10^-15^), pupal mass (F[2,376]=13.1, p<10^-5^), time to pupation (F[2,381]=35.9, p<10^-14^), and time to eclosion (F[2,345]=45.8, p<10^-15^) differ significantly between 1999 and 2024 (Fig S8). Survival to pupation during the recent studies (June 2024: 84%; July 2024: 57%) was higher than during the initial study (38%, Table 2).

## Discussion

Our findings demonstrate the potential for rapid evolution of thermal sensitivity in this population. The findings build on past evidence of rapid divergence in the temperature or photoperiod sensitivity of growth and developmental traits across North American populations of *P. rapae* (20–22) and other butterflies (23, 24). They also build on past observations of evolution in response to recent climate change in other pierid butterflies (25–27) to examine the underlying selection and evolutionary constraints. We closely followed the 1999 methodology, including collection and field study locations, rearing conditions and personnel, to isolate climate change as the primary difference between the 2024 and 1999 studies. Some coefficients and significance levels of the 1999 findings differ quantitively but not qualitatively from those previously reported (17, 18) due to loss of a small proportion of the initial data.

Multiple generations per year provide ample generations (∼100) for evolution to have occurred since the initial study. Seasonal plasticity in traits such as wing melanism has likely buffered variation in selection on butterflies across seasons that can slow evolution (28), but seasonal plasticity is unlikely to substantially influence the temperature sensitivity of caterpillar growth rate. Substantial gene flow among *P. rapae* populations in this region may have alternately enabled (via maintenance of genetic variation) or hindered (gene swamping) evolutionary responses (16). Western North American populations are genetically distinct from eastern North American populations and were likely founded by a small number of individuals (16). However, the populations maintain substantial genetic variation (heterozygosity) on which evolution can act (16).

Changing climates can drive complex temporal dynamics of selection and evolution. A previous resurvey for *Colias* butterflies suggested past selection for darker (more solar absorptive) wings to expand thermal opportunity daily and seasonally but a subsequent reversal in the direction of selection to avoid overheating with continued climate change (29). An analysis of our focal population suggests that shifts to higher thermal optima would increase growth in current thermal conditions but that an increasing incidence of warm temperatures over recent decades has reduced growth rates (28). More frequent warm temperatures over time may have intensified the previously documented selection (17, 18).

This increased capacity for growth at intermediate temperatures appears to have come at the cost of performance at the warmest temperatures (Figs 2, 4). As hot temperatures have become more common, selection has reversed to favor increased growth rate at the hottest temperatures. Selection in the system appears to continue to primarily act via temporally aggregated growth responses (quantified as pupal mass). As climate change proceeds, we expect selection on survival in response to thermal extremes to strengthen (30). We now detect significant selection via survival to grow faster at 29°C but the continuance of a non-significant tendency for lower growth rates at the highest temperature (35°C) to improve survival. This points to the importance of considering both environmental exposure and organismal sensitivity in anticipating evolutionary responses of thermal sensitivity (14).

Past quantitative genetic studies comparing full-sib families of our focal population indicated substantial genetic variation in growth rate, which increased more than thirty fold from low (8- 11°C) to high (35-40°C) temperatures (19). The past studies did not detect a genetic tradeoff between growth at 29°C and 35°C (which would be consistent with our 2024 phenotypic observation here). However, they did show that growth rates at 35°C and 40°C were strongly negatively correlated genetically (genetic correlation: -0.59) (19). Ongoing work is revisiting the quantitative genetic studies to confirm the evolution of thermal sensitivity and assess genetic constraints on the evolution of thermal sensitivity (31). Our resurvey suggests substantial increases in phenotypic variance of growth rates at 29 and 35°C and in negative correlations and covariances between growth rates at 29 and 35°C. The greater variation at higher temperatures increases the opportunity for selection, and potentially more rapid evolutionary responses, as the frequency of higher temperatures continues to increase with climate change (32, 33).

The complex temporal dynamics of selection and role of evolutionary constraints illustrates how repeating past studies of evolution and underlying selection can help assess the potential for evolutionary adaptation in response to climate change (6). Our resurvey suggests that phenotypic variance and evolutionary trade-offs will shape evolutionary responses to climate change.

Thermal opportunity and the risk of thermal stress will shift over time with climate change. Many evolutionary responses to climate change will likely entail complex temporal dynamics of selection and evolutionary response, such as those documented here. Past evolutionary responses to thermal opportunity may be reducing the ability of *P. rapae* to cope with hot temperatures that are increasing in incidences due to evolutionary tradeoffs. Shifts in selection and evolution through time may have implications for the demography and impacts of our focal global agricultural pest and we expect similar temporal dynamics for sensitive butterfly species.

## Materials and Methods

### Study organisms and rearing

*Pieris rapae* L. is native to Europe and is now widespread across much of North America, with multiple (3–5) generations per year in the Pacific Northwest. It is an agricultural pest on domesticated *Brassica* (16). *P. rapae* have five larval instars (over 90% of larval growth occurs in the final two instars) and have a facultative diapause (under photoperiod control) in the pupal stage. Caterpillars do not appear to actively regulate body temperatures (34, 35). More rapid caterpillar growth and development rates can result in greater survival to pupation, by decreasing time caterpillars exposed to enemies, including insect parasitoids and social wasp predators (36, 37). In addition, pupal mass is directly correlated with adult fecundity (34).

Studies in 2023 and 2024 closely followed those in 1999 including locations in Seattle, WA for field collection and the experimental garden and rearing conditions. Gravid females were caught in organic farms and community gardens and set up in individual cages in the UW Greenhouse with collard plants and nectar. Females were given 48 hours to lay eggs. Plants were watered daily and observed for egg hatching. 1st instar caterpillars were moved in groups of 15-20 caterpillars by family to a large petri dish with a collard leaf and a sponge to prevent the leaf from wilting. Caterpillars were reared in Argus environmental chambers (Conviron Argus Controls, Model No. G1000, Winnipeg, CA) with diurnal temperatures following a sawtooth pattern between 10 and 30°C and photoperiods of 16L:8D.

### Experimental garden

We planted an experimental garden of Champion Collard plants (C0350, Territorial Seed Co.) at the UW Farm at the Center of Urban Horticulture (47°39’31”N 122°17’30”W) to relate caterpillar temperature sensitivity of growth to the following field outcomes: survival to pupation, time to pupation, pupal mass (mg), and (for a subset of caterpillars) egg production. The garden consisted of 11 rows, 1 m apart and 15 m in length. The experimental garden housed 260 greenhouse raised plants inserted in holes in black weed cloth at 0.75 m intervals. Plants were drip irrigated daily. Wasp traps were placed around the perimeter of the garden to limit predation.

We recorded operative temperatures at 5minute intervals throughout the summer using caterpillar models [type T thermocouples (OMEGA Wire Inc PP-T-24-SLE-30M) imbedded in mechanically set epoxy painted leaf green] that were placed on the underside of the outer leaves of plants distributed throughout the garden. In 1999, twenty caterpillar models were monitored using a Campbell 20XL datalogger. In 2024, twelve models were monitored using HOBO data loggers (Onset Computer Company UX120-014). Long term daily weather station maximum and minimum temperatures were downloaded from the Global Historical Climatology Network daily (GHCNd) for a station near the experimental garden (USW00094290). We estimated summer hourly temperatures during an initial (1990–1999) and recent period (2015–2024) using the R function diurnal_temp_variation_sine in the TrenchR package (38).

The timing of the 2024 experiments accounted for seasonal shifts in environmental temperature and phenology since the initial studies (Fig 1a). The August 1999 experiment was conducted between August 11-25, with most caterpillars moved to the garden on August 12-13 (n=206 from 20 families). For the June 2024 experiment, batches of caterpillars were moved into the garden on June 22 and 23 (n=161 from 7 families). For the July 2024 experiment, batches of caterpillars were moved into the garden on July 28 and 29 (n=303 from 23 families). To assess alternative explanations for temporal shifts such as parental effects, we additionally analyzed growth rate data for assays that initiated on July 8 (n=214 from 16 families) and 30 (n=139 from 15 families), 2023 (Figure S8). We omit analyses of the corresponding garden data due to high mortality, primarily from wasp predation.

### Thermal sensitivity

Newly molted 4th instar caterpillars were selected by family, placed individually in petri dishes, and weighed using a Mettler Toledo scale (Model XSR105DU). Caterpillars over 20mg were disqualified to prevent pupation during the experiment. We measured short-term relative growth rates (g/g/hr) feeding on small collard leaves sequentially in the following treatments: 23°C (7 hours), 11°C (15 hours overnight), 29°C (5 hours), 35°C (4 hours), and 17°C (15 hours overnight).

### Selection experiment

Following the last morning weighing, petri dishes were shuffled, and caterpillars were transported to the experiment garden, where they were placed randomly and individually in the inner whorl of the collard plants. To prevent predation, plants were covered in bridal veil either by row (1999) or individually with twist ties (2024). Caterpillars were censused each morning between 8:00 am and 12:00 pm until all surviving caterpillars had pupated. Retrieved pupae were kept in 16 oz clear SOLO cups with a piece of mesh at room temperature and monitored daily for eclosing adults.

Pupae were weighed after 48 hours of hardening. Eclosing adults were sexed and weighed. To assess fecundity for the August 1999 and July 2024 experiment, recent eclosed females were marked with a series of dots on their forewing in a cold room (4°C). They were allowed to mate with males with different mothers for 48 hours in a large cage. Eggs were counted 48 hours after females were moved to individual cages with nectar and a collard plant.

### Analyses

We used linear models and ANOVAs (R functions lm and anova) to analyze shifts through time along with linear-mixed-effects models (R function lme in package nlme) accounting for mother as a random effect. For TPCs, temperature was included as a third-degree polynomial. We used generalized linear models with the binomial family to model survival as a binary response variable. We used Chi tests to compare models of fitness responses with and without interactions and report the model with more support. To assess shifts in distributions through time, we used Levene tests to assess changes in variance followed by Welch tests or t-tests for the cases of unequal and equal variances, respectively.

We modelled growth rates at each temperature as individual traits. Models of fitness responses (pupal mass, survival, and development time) accounted for initial mass. We centered the values of growth rate traits by subtracting means. We estimated selection gradients using linear multiple regressions standardizing trait values and fitness responses by dividing by means. We estimated phenotypic variance covariance (P matrices, R function var) and correlation (R function cor) matrices. We estimated eigenvectors (R function eigen) to describe the primary components of variation in growth rates across temperatures. We compared variance covariance matrices across time using random skewers (R function RandomSkewers in the evolqg package) (39, 40). We compared correlation matrices across time using Mantel permutation tests (R function MantelCor in the evolqg package).

## Data, Materials, and Software Availability

Data and code are available at https://github.com/HuckleyLab/PrapaeGardenExpt and will be archived in Zenodo.

## Acknowledgments

We would like to thank and acknowledge the student volunteers from Edmonds Community College who assisted in data collection, along with University of Washington (UW) undergraduate researchers. Thank you to Jared Haar, Jennifer Lopez, Lucie Reizian, Marcos Alvarez, Samuel Moles, Max Oberholtzer, Anna Brasket, Alex Rieflin, and Hannah Woods. We thank the UW Farm and the UW Biology Greenhouse for their assistance. Amy Angert, Louie Yang, and members of our research groups provided constructive comments on the manuscript. This work was supported by the National Science Foundation (IOS-2222089 to L.B.B., IOS-2222090 to J.G.K.).

## Supporting information

**Figure S1.**
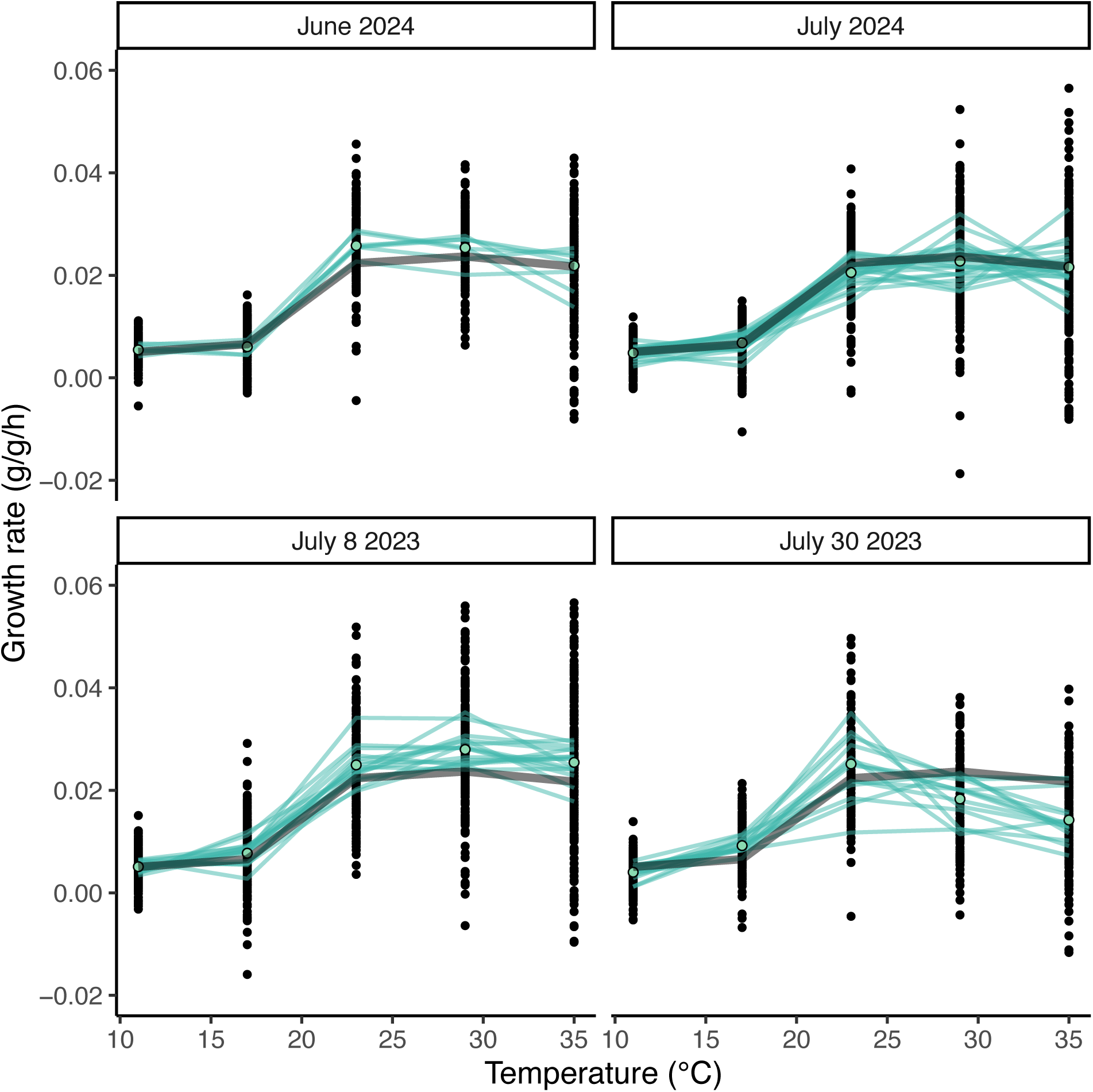
Across recent seasons and years, the thermal sensitivity of caterpillar relative growth rates (RGR, g/g/h) shows declines at the highest measurement temperatures. We depict individual values and means ± SE at each measurement temperature along with means by family and for the 2024 selection studies.

**Figure S2.**
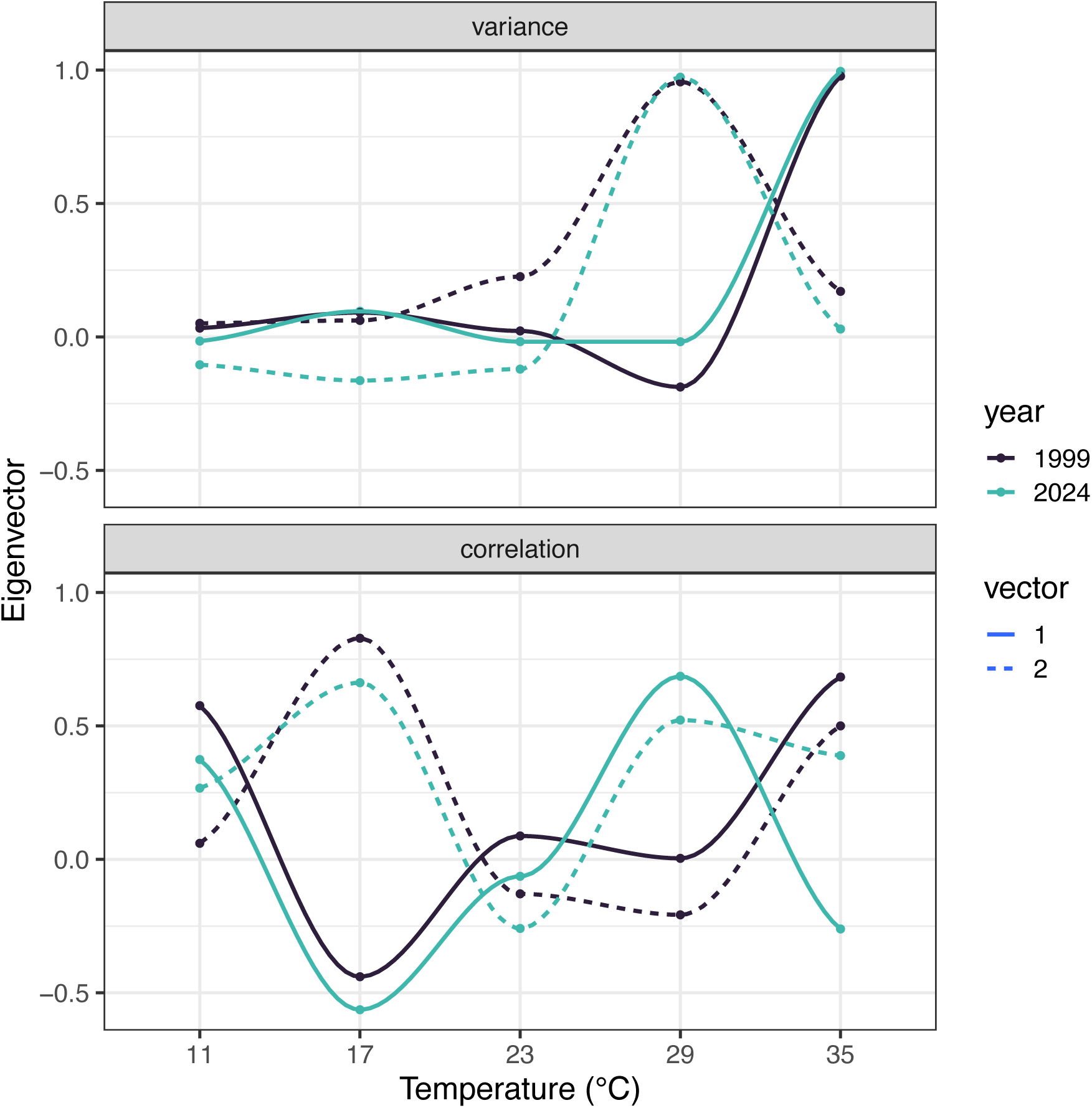
The first and second eigenvectors corresponding to the variance covariance matrix (accounting for 38% and 35% of past variation and 54% and 22% of recent variation, respectively) illustrate that the variance structure has remained similar over time. The first and second eigenvectors corresponding to the correlation matrix (accounting for 29% and 23% of past variation and 29% and 25% of recent variation, respectively) indicate weakening correlations at the extreme temperatures over time.

**Figure S3.**
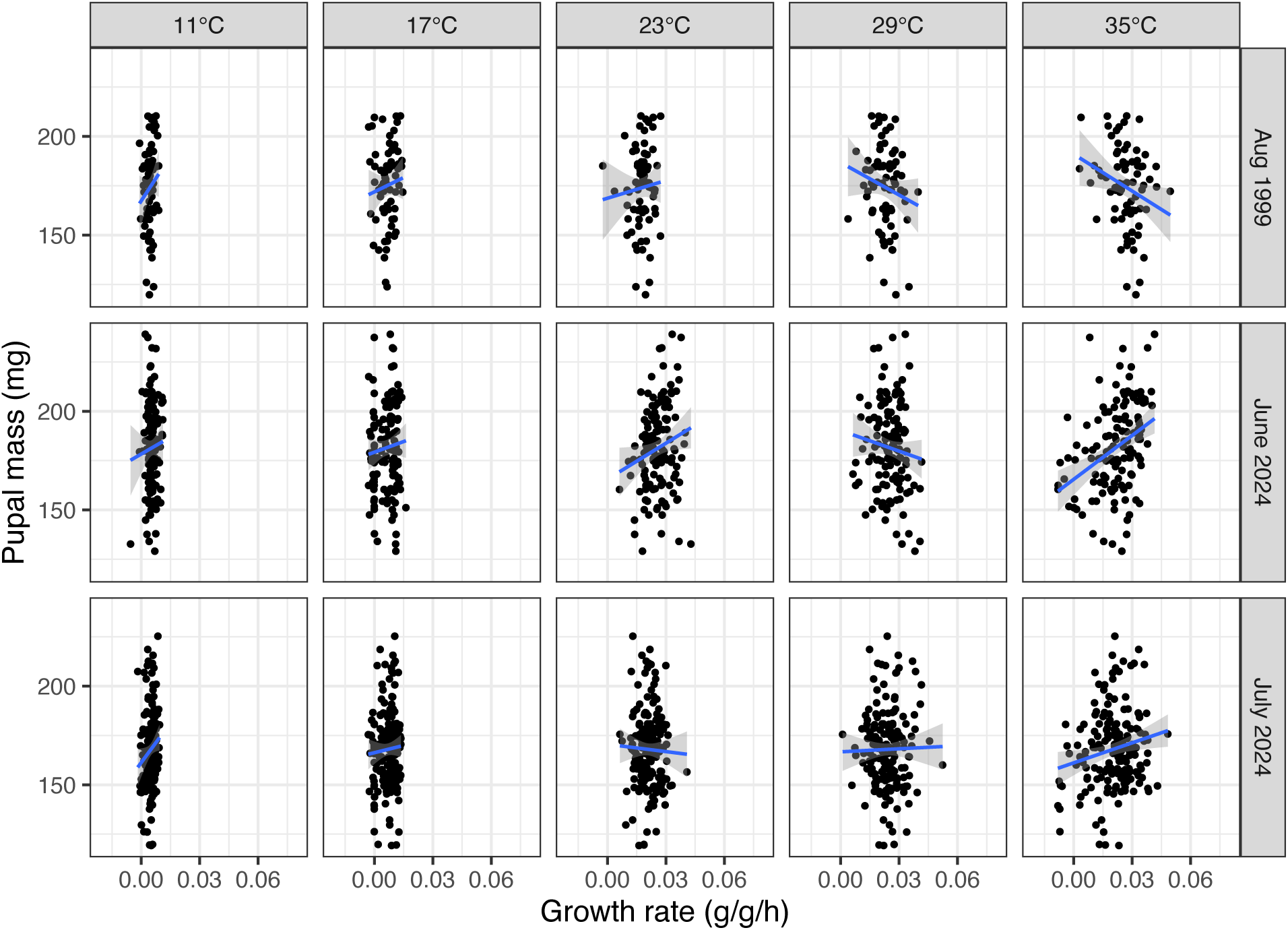
Selection on growth rates via pupal mass. We depict linear regressions for growth rate at each temperature, which differs from our estimation of selection gradients in the main text.

**Figure S4.**
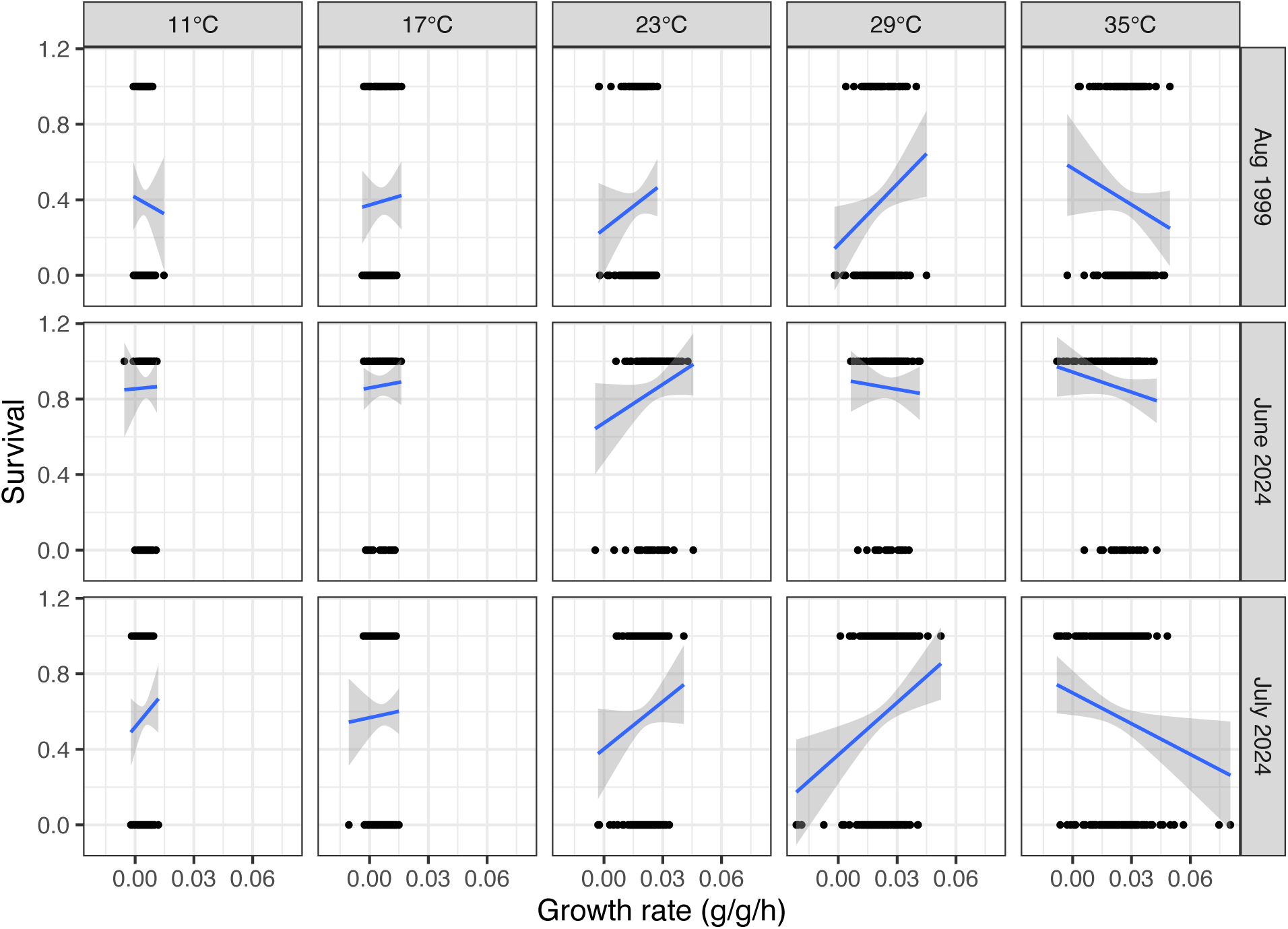
Selection on growth rates via survival. We depict linear regressions for growth rate at each temperature, which differs from our estimation of selection gradients in the main text.

**Figure S5.**
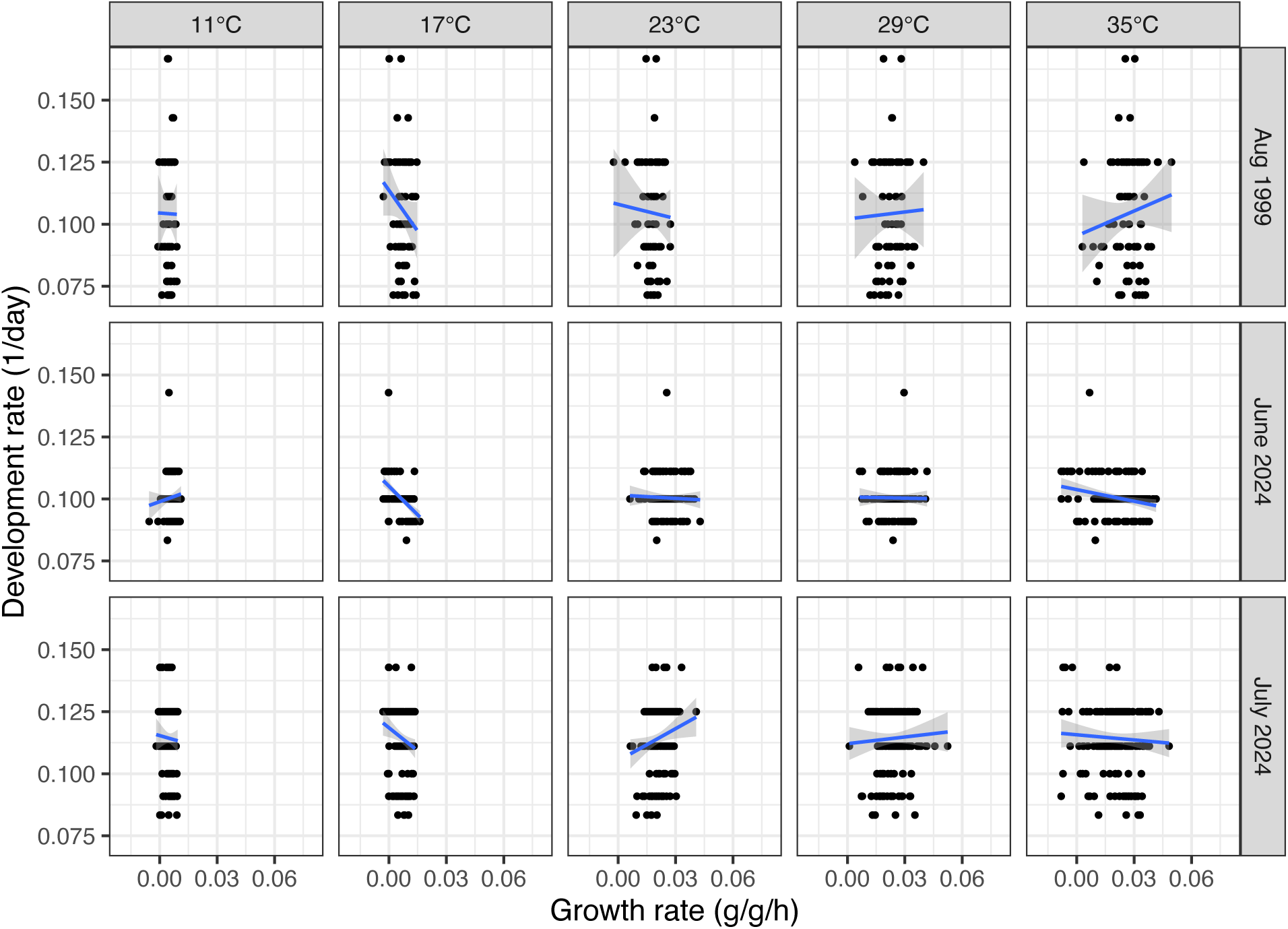
Selection on growth rates via development rate. We depict linear regressions for growth rate at each temperature, which differs from our estimation of selection gradients in the main text.

**Figure S6.**
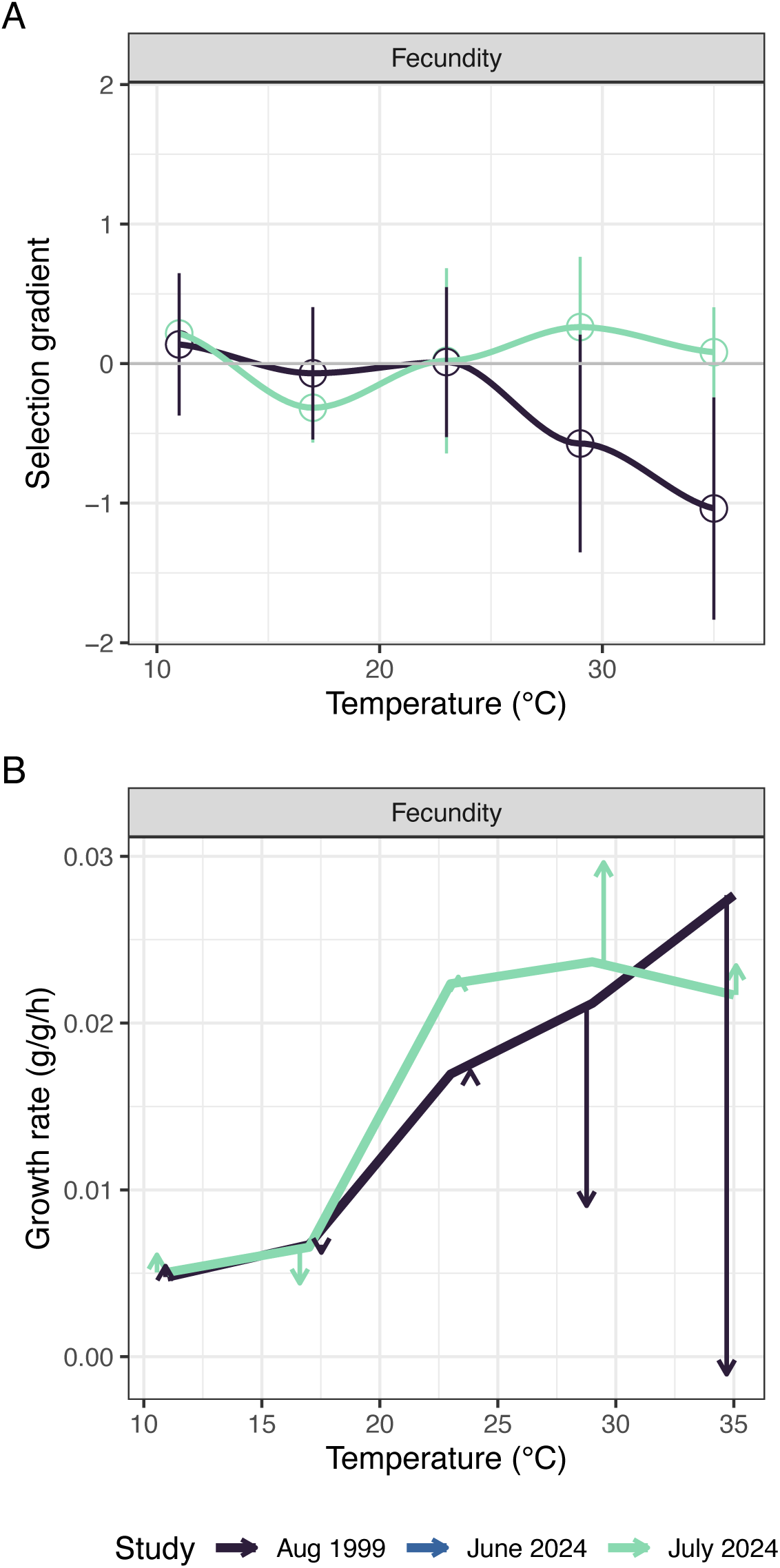
A) Estimated selection gradients (±SE) for fecundity have changed between the past and recent studies. Selection gradients that are statistically significant are shaded. B) We overlay the selection gradients (arrow lengths indicate magnitudes) to illustrate potential shifts in thermal sensitivity.

**Figure S7.**
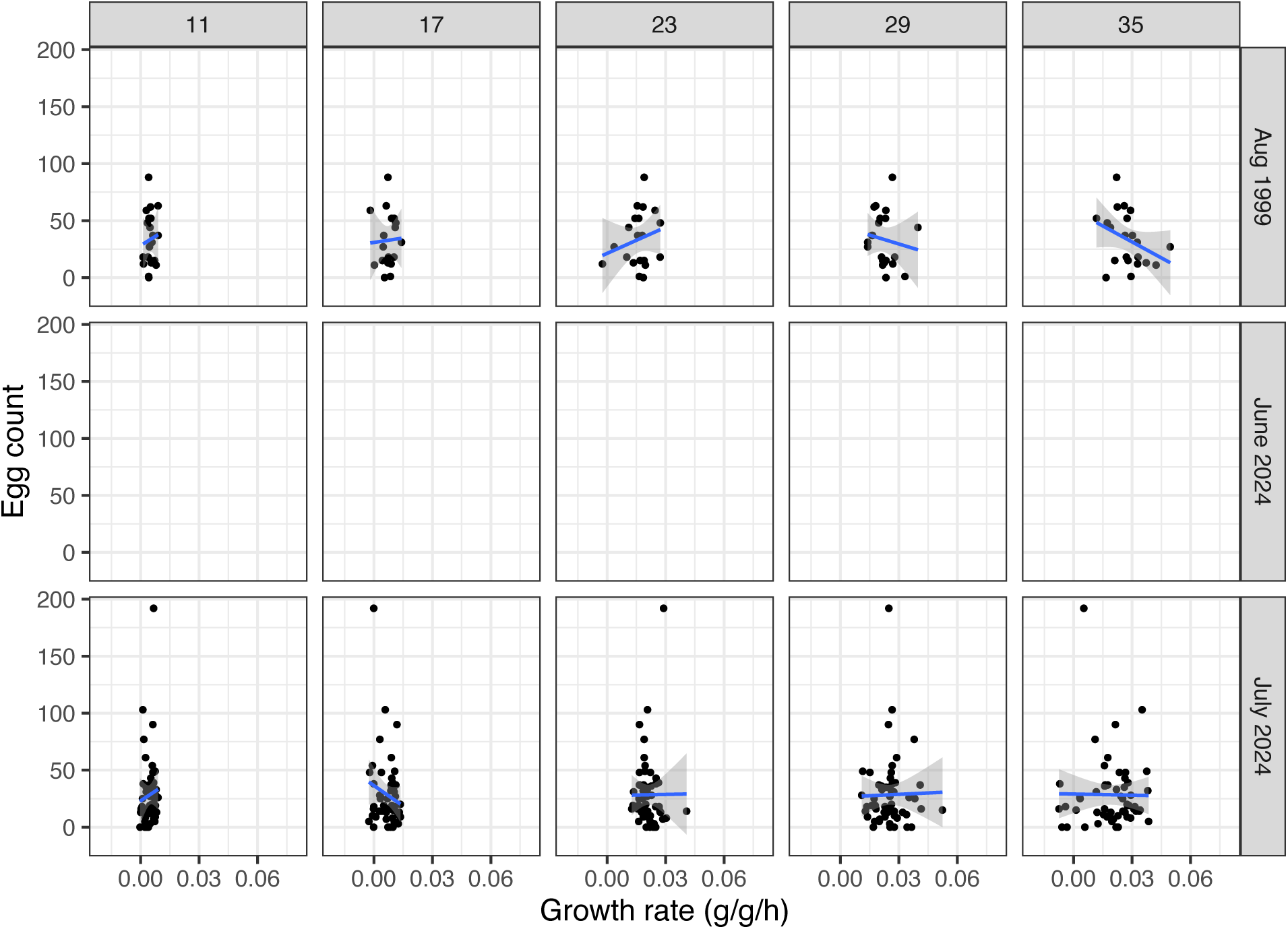
Selection on growth rates via fecundity. We depict linear regressions for growth rate at each temperature, which differs from our estimation of selection gradients in the main text.

**Figure S8.**
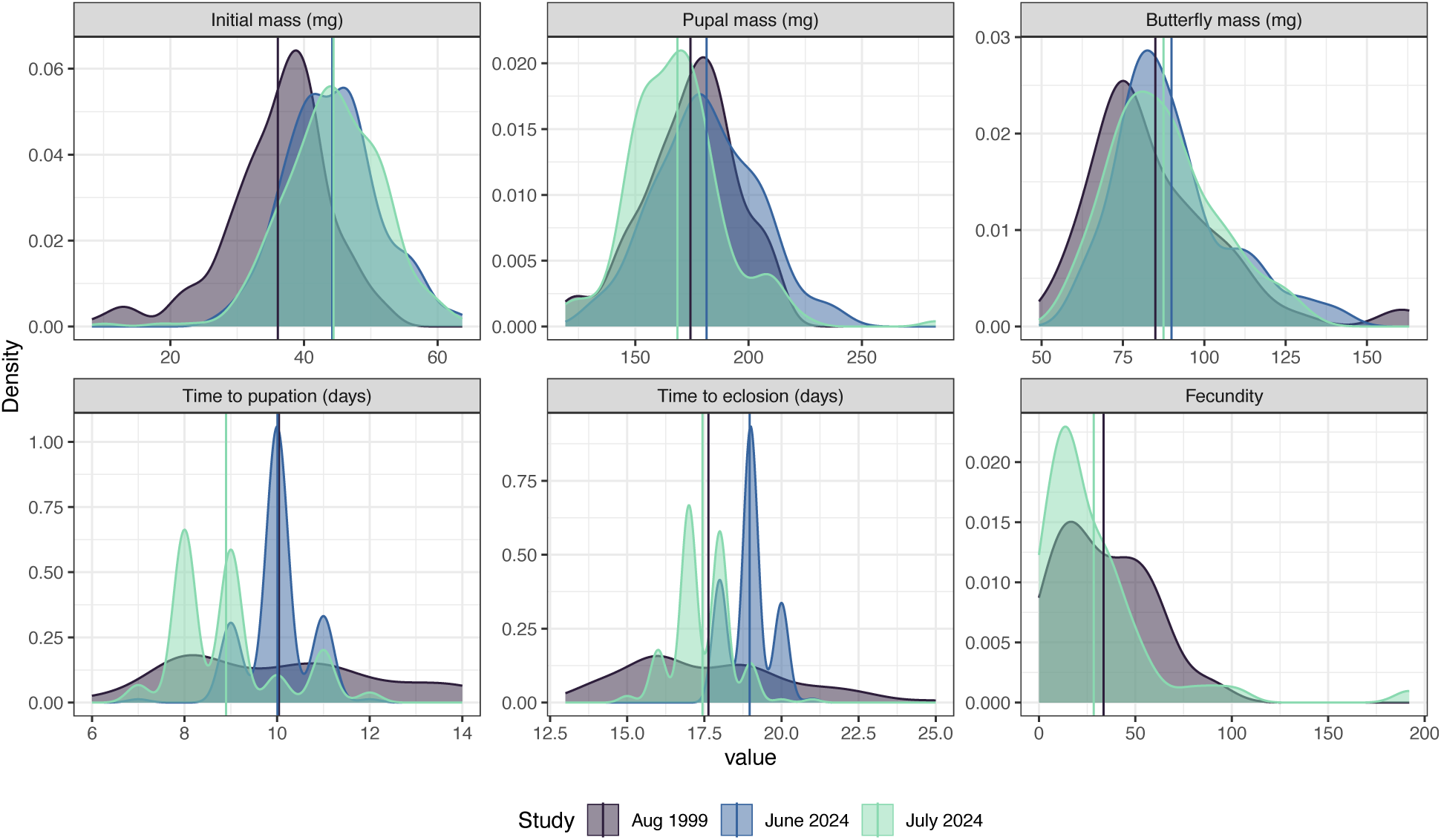
The distributions of traits and fitness responses through the studies.

**Table S1.**
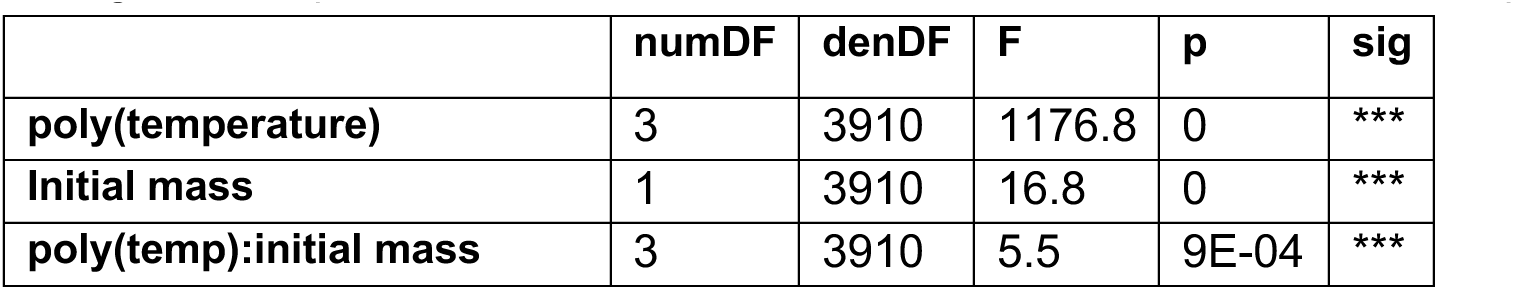
ANOVA results from a linear mixed effects model with a random effect of mother nested within the experiments in Extended Data Figure 1. The standard deviations of the random effects (sigma: overall=0.0085, experiment=0.0026, mother=0.0006) indicate that differences among recent experiments are smaller than those between the 1999 and 2024 experiments.

**Table S2.** Variance covariance (P) and correlation matrices comparing relative growth rates (RGR) across temperatures.

|  | RGR11 | RGR17 | RGR23 | RGR29 | RGR35 | type | period |
| --- | --- | --- | --- | --- | --- | --- | --- |
| RGR11 | 5.812 | -0.785 | -1.643 | -5.389 | 0.258 | P | recent |
| RGR17 | -0.785 | 21.493 | -1.936 | 1.527 | 14.779 | P | recent |
| RGR23 | -1.643 | -1.936 | 42.17 | 4.15 | -1.367 | P | recent |
| RGR29 | -5.389 | 1.527 | 4.15 | 69.902 | -30.81 | P | recent |
| RGR35 | 0.258 | 14.779 | -1.367 | -30.81 | 133.385 | P | recent |
| RGR111 | 5.587 | -1.73 | -2.433 | -5.075 | -0.001 | P | initial |
| RGR171 | -1.73 | 17.575 | -2.984 | -2.862 | 1.295 | P | initial |
| RGR231 | -2.433 | -2.984 | 28.085 | 8.545 | 0.525 | P | initial |
| RGR291 | -5.075 | -2.862 | 8.545 | 57.798 | -1.858 | P | initial |
| RGR351 | -0.001 | 1.295 | 0.525 | -1.858 | 64.36 | P | initial |
| RGR112 | 1 | -0.07 | -0.105 | -0.267 | 0.009 | correlation | recent |
| RGR172 | -0.07 | 1 | -0.064 | 0.039 | 0.276 | correlation | recent |
| RGR232 | -0.105 | -0.064 | 1 | 0.076 | -0.018 | correlation | recent |
| RGR292 | -0.267 | 0.039 | 0.076 | 1 | -0.319 | correlation | recent |
| RGR352 | 0.009 | 0.276 | -0.018 | -0.319 | 1 | correlation | recent |
| RGR113 | 1 | -0.175 | -0.194 | -0.282 | 0 | correlation | initial |
| RGR173 | -0.175 | 1 | -0.134 | -0.09 | 0.038 | correlation | initial |
| RGR233 | -0.194 | -0.134 | 1 | 0.212 | 0.012 | correlation | initial |
| RGR293 | -0.282 | -0.09 | 0.212 | 1 | -0.03 | correlation | initial |
| RGR353 | 0 | 0.038 | 0.012 | -0.03 | 1 | correlation | initial |

**Table S3.**
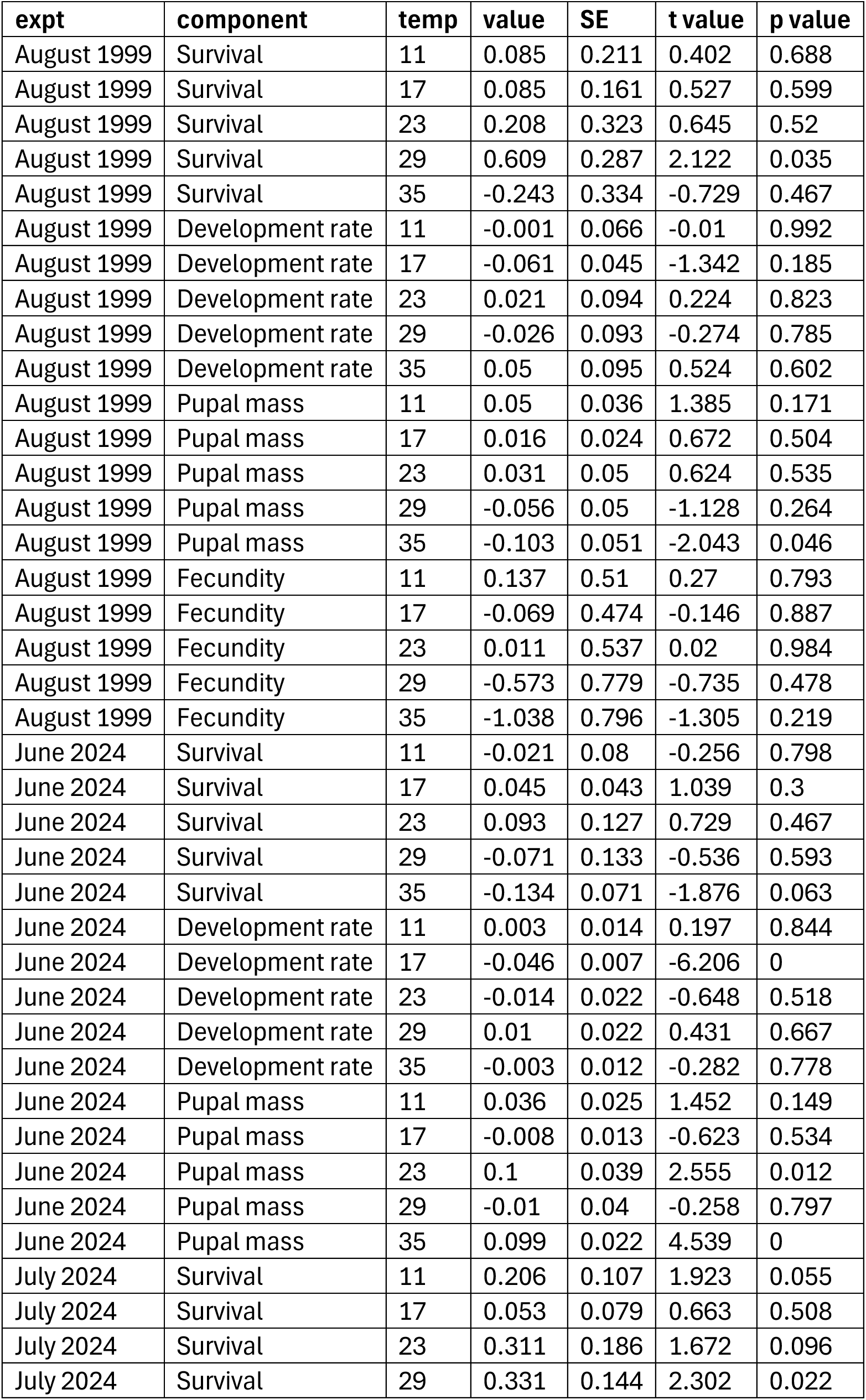
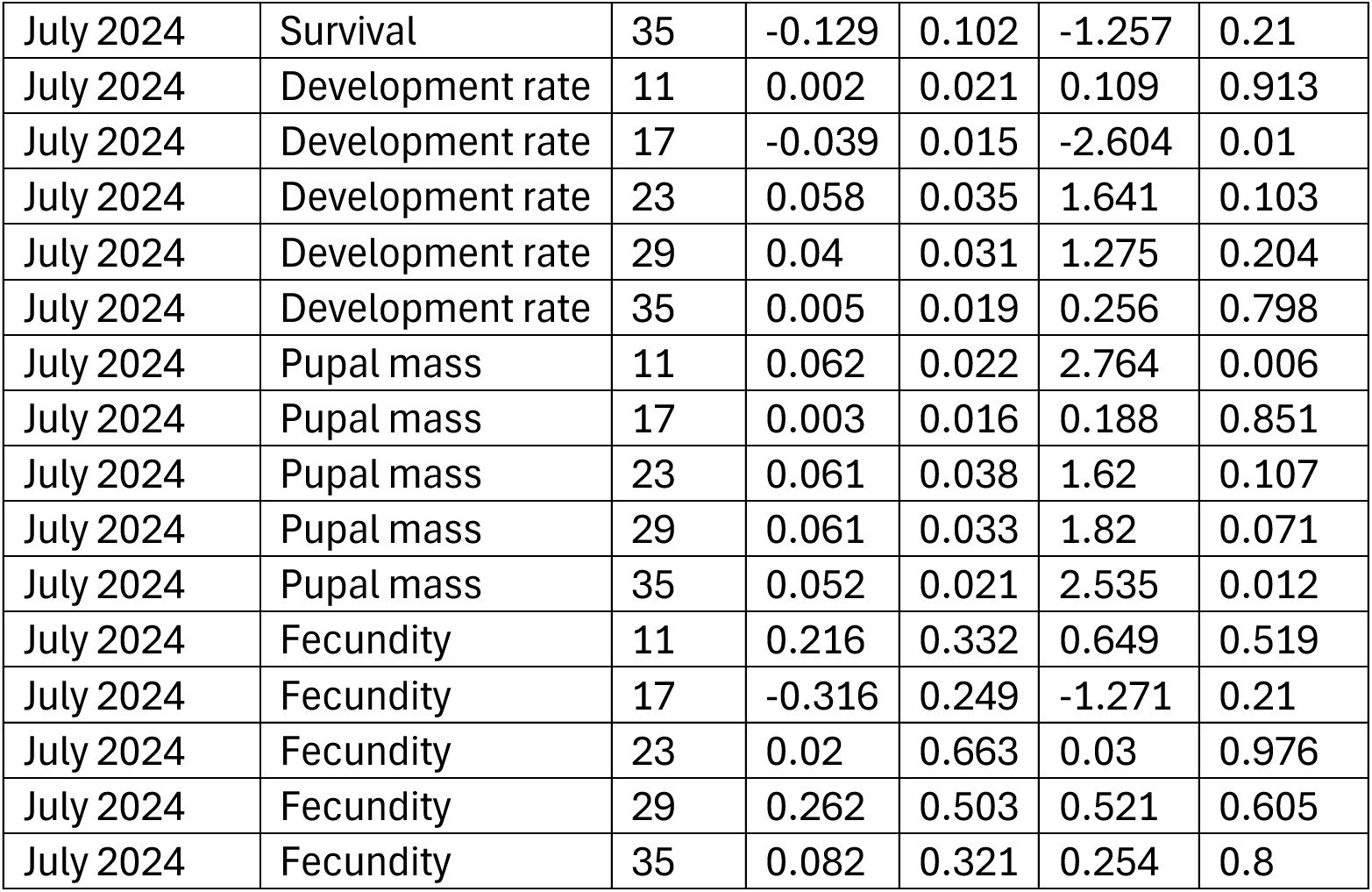
Mean-standardized selection coefficients (±SE) along with t and p values across the experiments and fitness components.

| expt | component | temp | value | SE | t value | p value |
| --- | --- | --- | --- | --- | --- | --- |
| August 1999 | Survival | 11 | 0.085 | 0.211 | 0.402 | 0.688 |
| August 1999 | Survival | 17 | 0.085 | 0.161 | 0.527 | 0.599 |
| August 1999 | Survival | 23 | 0.208 | 0.323 | 0.645 | 0.52 |
| August 1999 | Survival | 29 | 0.609 | 0.287 | 2.122 | 0.035 |
| August 1999 | Survival | 35 | -0.243 | 0.334 | -0.729 | 0.467 |
| August 1999 | Development rate | 11 | -0.001 | 0.066 | -0.01 | 0.992 |
| August 1999 | Development rate | 17 | -0.061 | 0.045 | -1.342 | 0.185 |
| August 1999 | Development rate | 23 | 0.021 | 0.094 | 0.224 | 0.823 |
| August 1999 | Development rate | 29 | -0.026 | 0.093 | -0.274 | 0.785 |
| August 1999 | Development rate | 35 | 0.05 | 0.095 | 0.524 | 0.602 |
| August 1999 | Pupal mass | 11 | 0.05 | 0.036 | 1.385 | 0.171 |
| August 1999 | Pupal mass | 17 | 0.016 | 0.024 | 0.672 | 0.504 |
| August 1999 | Pupal mass | 23 | 0.031 | 0.05 | 0.624 | 0.535 |
| August 1999 | Pupal mass | 29 | -0.056 | 0.05 | -1.128 | 0.264 |
| August 1999 | Pupal mass | 35 | -0.103 | 0.051 | -2.043 | 0.046 |
| August 1999 | Fecundity | 11 | 0.137 | 0.51 | 0.27 | 0.793 |
| August 1999 | Fecundity | 17 | -0.069 | 0.474 | -0.146 | 0.887 |
| August 1999 | Fecundity | 23 | 0.011 | 0.537 | 0.02 | 0.984 |
| August 1999 | Fecundity | 29 | -0.573 | 0.779 | -0.735 | 0.478 |
| August 1999 | Fecundity | 35 | -1.038 | 0.796 | -1.305 | 0.219 |
| June 2024 | Survival | 11 | -0.021 | 0.08 | -0.256 | 0.798 |
| June 2024 | Survival | 17 | 0.045 | 0.043 | 1.039 | 0.3 |
| June 2024 | Survival | 23 | 0.093 | 0.127 | 0.729 | 0.467 |
| June 2024 | Survival | 29 | -0.071 | 0.133 | -0.536 | 0.593 |
| June 2024 | Survival | 35 | -0.134 | 0.071 | -1.876 | 0.063 |
| June 2024 | Development rate | 11 | 0.003 | 0.014 | 0.197 | 0.844 |
| June 2024 | Development rate | 17 | -0.046 | 0.007 | -6.206 | 0 |
| June 2024 | Development rate | 23 | -0.014 | 0.022 | -0.648 | 0.518 |
| June 2024 | Development rate | 29 | 0.01 | 0.022 | 0.431 | 0.667 |
| June 2024 | Development rate | 35 | -0.003 | 0.012 | -0.282 | 0.778 |
| June 2024 | Pupal mass | 11 | 0.036 | 0.025 | 1.452 | 0.149 |
| June 2024 | Pupal mass | 17 | -0.008 | 0.013 | -0.623 | 0.534 |
| June 2024 | Pupal mass | 23 | 0.1 | 0.039 | 2.555 | 0.012 |
| June 2024 | Pupal mass | 29 | -0.01 | 0.04 | -0.258 | 0.797 |
| June 2024 | Pupal mass | 35 | 0.099 | 0.022 | 4.539 | 0 |
| July 2024 | Survival | 11 | 0.206 | 0.107 | 1.923 | 0.055 |
| July 2024 | Survival | 17 | 0.053 | 0.079 | 0.663 | 0.508 |
| July 2024 | Survival | 23 | 0.311 | 0.186 | 1.672 | 0.096 |
| July 2024 | Survival | 29 | 0.331 | 0.144 | 2.302 | 0.022 |
| July 2024 | Survival | 35 | -0.129 | 0.102 | -1.257 | 0.21 |
| July 2024 | Development rate | 11 | 0.002 | 0.021 | 0.109 | 0.913 |
| July 2024 | Development rate | 17 | -0.039 | 0.015 | -2.604 | 0.01 |
| July 2024 | Development rate | 23 | 0.058 | 0.035 | 1.641 | 0.103 |
| July 2024 | Development rate | 29 | 0.04 | 0.031 | 1.275 | 0.204 |
| July 2024 | Development rate | 35 | 0.005 | 0.019 | 0.256 | 0.798 |
| July 2024 | Pupal mass | 11 | 0.062 | 0.022 | 2.764 | 0.006 |
| July 2024 | Pupal mass | 17 | 0.003 | 0.016 | 0.188 | 0.851 |
| July 2024 | Pupal mass | 23 | 0.061 | 0.038 | 1.62 | 0.107 |
| July 2024 | Pupal mass | 29 | 0.061 | 0.033 | 1.82 | 0.071 |
| July 2024 | Pupal mass | 35 | 0.052 | 0.021 | 2.535 | 0.012 |
| July 2024 | Fecundity | 11 | 0.216 | 0.332 | 0.649 | 0.519 |
| July 2024 | Fecundity | 17 | -0.316 | 0.249 | -1.271 | 0.21 |
| July 2024 | Fecundity | 23 | 0.02 | 0.663 | 0.03 | 0.976 |
| July 2024 | Fecundity | 29 | 0.262 | 0.503 | 0.521 | 0.605 |
| July 2024 | Fecundity | 35 | 0.082 | 0.321 | 0.254 | 0.8 |

**Table S4.** The mean and standard deviation (SD) of fitness outcomes for each experiment.

| **Experiment** | **Survival** | **SD** | **Dev Rate** | **SD** | **Pupal mass** | **SD** | **Fecundity** | **SD** |
| --- | --- | --- | --- | --- | --- | --- | --- | --- |
| August 1999 | 0.383 | 0.487 | 0.1042 | 0.0224 | 174.33 | 20.68 | 33.48 | 23.57 |
| June 2024 | 0.839 | 0.369 | 0.1004 | 0.0073 | 181.45 | 22.41 | NA | NA |
| July 2024 | 0.571 | 0.496 | 0.1141 | 0.0136 | 168.6 | 21.6 | 28.41 | 31.56 |

## Notes

### Competing Interest Statement

The authors have declared no competing interest.

